# The world’s largest High Arctic lake is dominated by uncharacterized, genetically highly diverse phages

**DOI:** 10.1101/2024.09.10.612304

**Authors:** Audrée Lemieux, Alexandre J. Poulain, Stéphane Aris-Brosou

## Abstract

While the Earth’s virosphere is estimated to be in the range of 10^31^ viral particles, the vast majority of its diversity has yet to be discovered. In recent years, metagenomics has rapidly allowed the identification of viruses, from microenvironments to extreme environments like the High Arctic. However, the High Arctic virome is largely composed of viral sequences that have few matches to classified viruses in existing databases. Here, to bypass limitations posed by similarity-based strategies, we resorted to a paired metagenomic (DNA, *n* = 8) and metatranscriptomic (RNA, *n* = 6) approach that placed viral genes found in Lake Hazen, the world’s largest High Arctic lake, in a phylogenetic context with known viruses. We show that High Arctic viruses have undergone unique evolutionary processes characterized by high evolutionary rates, making them distinct from and more diverse than known viruses.

## Introduction

Viruses are estimated to number on the order of 10^31^ particles on Earth (Mushegian, 2020), occupying an extraordinary range of habitats from localized microenvironments to extreme ecosystems. Yet, despite this vast abundance, only 17,554 viral species have been formally classified by the International Committee on Taxonomy of Viruses (as of March 20, 2026) (International Committee on Taxonomy of Viruses, 2026). Viral research has historically concentrated on pathogens of humans, plants, and animals (Gil et al., 2021), leaving much of the global virosphere, particularly in extreme and remote environments, largely unexplored.

In previous work (Lemieux, Colby, Poulain, & Aris-Brosou, 2022), we characterized viruses and their eukaryotic hosts from soils and sediments collected at Lake Hazen, the world’s largest High Arctic freshwater lake (Lehnherr et al., 2018), using similarity-based approaches. Strikingly, known viruses accounted for less than 1% of recovered sequences, despite estimates of approximately 10^9^ viral particles per gram of soil or sediment (Dávila-Ramos et al., 2019). Moreover, more than 50% of sequences could not be assigned to any superkingdom (Lemieux et al., 2022). These observations suggest that a substantial fraction of viral diversity remains uncharacterized, likely reflecting the limitations of homology-based methods when applied to novel viruses from poorly sampled environments (Dávila-Ramos et al., 2019).

Relatively few studies have attempted to comprehensively assess viral diversity in Arctic ecosystems (Yau & Seth-Pasricha, 2019), and those that have consistently report viromes dominated by unknown sequences. Most investigations have focused on DNA viruses (Aguirre de Cárcer, López-Bueno, Pearce, & Alcamí, 2015; Emerson et al., 2018; Zhong et al., 2020, 2024; Wang et al., 2022; Kim et al., 2024; Angly et al., 2006; Calayag et al., 2025; Demina et al., 2025; Pettersson et al., 2025), while RNA viruses remain comparatively understudied due to their smaller genomes and the inherent instability of RNA (Yau & Seth-Pasricha, 2019). Studies reconstructing viral assembled genomes (VAGs) have further highlighted this knowledge gap, revealing limited similarity between Arctic viral genes and existing database sequences (Labbé, Girard, Vincent, & Culley, 2020; Trubl et al., 2021), as well as between recovered viral genomes and established references (Nguyen, Robertsen, & Landfald, 2017). These findings raise the fundamental question about how evolutionarily related are Arctic viral communities, particularly those in Lake Hazen, to currently characterized viruses.

To address this outstanding question, we combined metagenomic and metatranscriptomic approaches to place Lake Hazen viruses within a phylogenetic framework alongside reference viral sequences. By integrating predicted genes from our datasets with publicly available viral genomes, we show that the Arctic virome is highly divergent from known viruses, characterized by a predominance of bacteriophages and substantially greater genetic diversity than represented in current databases.

## Materials and Methods

### Sample collection, processing, and sequencing

High Arctic sediments and soil cores were collected from Lake Hazen (Nunavut, Canada) and processed as previously described (Lemieux et al., 2022; Colby et al., 2020). RNA (*n* = 6) and DNA (*n* = 8) metagenomic libraries were prepared and sequenced on an Illumina HiSeq 2500 platform (Illumina, San Diego, CA, USA) at Génome Québec using paired-end 125 bp reads. Each library was replicated two (DNA) or three (RNA) times. Raw reads were trimmed for adapters and low-quality bases using Trimmomatic v0.39 (Bolger, Lohse, & Usadel, 2014) with settings: `TruSeq3-PE.fa:2:30:10 LEADING:3 TRAILING:3 SLIDINGWINDOW:4:15 MINLEN:36`. Contigs were assembled with MEGAHIT v1.2.9 (D. Li, Liu, Luo, Sadakane, & Lam, 2015; D. Li et al., 2016) (default parameters). Contigs ≥300 bp from soil and sediment samples were pooled separately for RNA and DNA data to reconstruct viral genes independently (Figure 1).

**Figure 1.**
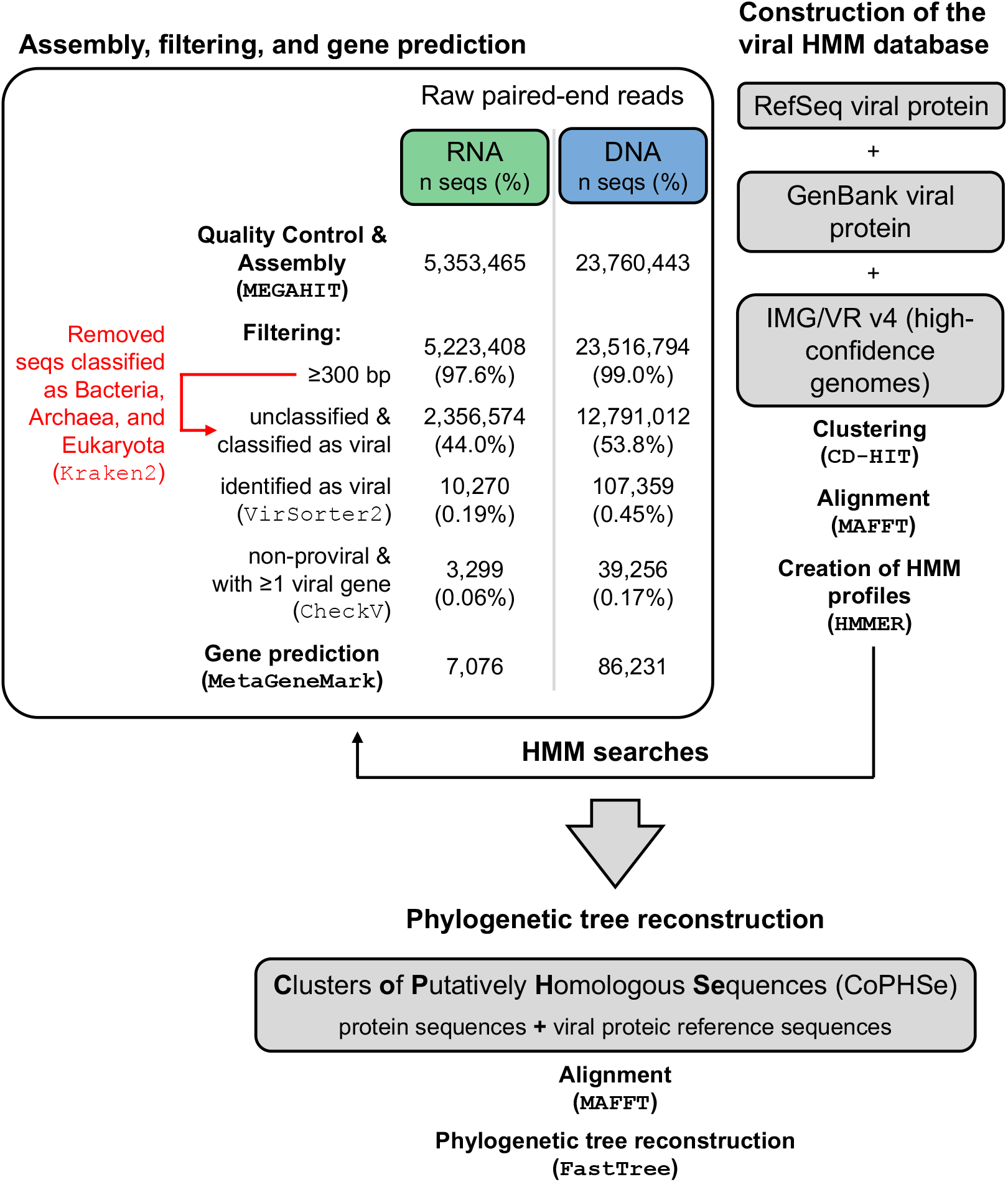
Analysis pipeline of the RNA and DNA data.

### Viral genome identification

Contigs were first filtered using Kraken2 v2.1.1 (Wood, Lu, & Langmead, 2019) against its Standard database (April 2, 2025; https://benlangmead.github.io/aws-indexes/k2), which includes archaeal, bacterial, viral, plasmid, human, and vector sequences. `The extract_kraken_reads.py` script from KrakenTools v1.2 (Lu, 2021) removed contigs classified as bacterial (taxID 2), archaeal (taxID 2157), or eukaryotic (taxID 2759) using parameters `-t 2 2157 2759 --include-children –exclude`. Unclassified contigs and those assigned to other taxa (including Viruses) were retained.

Viral RNA and DNA genomes were identified with VirSorter2 v2.2.4 (Guo et al., 2021) (default parameters). CheckV (Nayfach et al., 2021) (default parameters) assessed prediction quality, retaining non-proviral sequences with ≥1 viral gene at high (*>*90%), medium (50–90%), or low (*<*5%) completeness, excluding those with warnings (e.g., low-confidence DTRs or high k-mer duplication). Protein-coding genes were predicted using MetaGeneMark v3.38 (Besemer & Borodovsky, 1999; Zhu, Lomsadze, & Borodovsky, 2010) (default parameters).

### Clusters of Putatively Homologous Sequences (CoPHSe)

A viral reference database was built from 169.5 million protein sequences: RefSeq v230 (May 5, 2025; https://ftp.ncbi.nlm.nih.gov/refseq/release/viral/viral.*.protein.faa.gz), GenBank v266 (April 30, 2025; https://ftp.ncbi.nlm.nih.gov/ncbi-asn1/protein_fasta/gbvrl*.fsa_aa.gz), and IMG/VR v4.1 high-confidence genomes (Camargo et al., 2022). Influenza and SARS-CoV-2 sequences comprised 87.0% of RefSeq/GenBank entries.

Sequences were dereplicated and clustered at 90% identity using CD-HIT v4.8.1 (W. Li & Godzik, 2006; Fu, Niu, Zhu, Wu, & Li, 2012; Emerson et al., 2018), yielding 31,758,083 clusters. Clusters with 10–99 reference sequences were retained (*N* = 1, 910, 901; 6.0% of total; Supplementary Figure 1). Multiple sequence alignments were generated with MAFFT v7.471 (Katoh, Misawa, Kuma, & Miyata, 2002; Katoh & Standley, 2013) (`--retree 1`), and HMM profiles built with HMMER v3.2.1 (Eddy, 2011). Our predicted genes were clustered with references via HMM searches (*E*-value thresholds: 10^−15^, 10^−20^), termed **C**lusters **o**f **P**utatively **H**omologous **Se**quences (CoPHSe).

CoPHSe alignments (≥3 predicted + ≥1 reference sequences) were trimmed with tri-mAl v1.4 (Capella-Gutiérrez, Silla-Martínez, & Gabaldón, 2009) (gappyout mode) and trees built using FastTree v2.1.11 (Price, Dehal, & Arkin, 2009, 2010) under the LG+Γ model (Le & Gascuel, 2008). Phage and non-phage sequences were separated using IMG/VR taxonomy, ViralZone annotations (Hulo et al., 2010), and organism names/keywords from RefSeq/GenBank (manually verified). CoPHSe were categorized by reference consen-sus (100% agreement): (i) non-phage, (ii) phage, (iii) unknown, (iv) non-phage+phage, (v) non-phage+phage+unknown, (vi) non-phage+unknown, (vii) phage+unknown. Mixed categories (iv/v; *<*0.05%) were excluded.

### Phylogenetic and taxonomic assessment

In most cases, viral predicted genes could be grouped into a single monophyletic clade, which was used to reroot trees using the `getMRCA` and `root` functions from `ape` v5.8 (Paradis & Schliep, 2019). The relatedness and evolution of RNA and DNA viral genes were then assessed through the phylogenetic trees. First, to test if viruses from the High Arctic are more diverse than those from the viral reference database (mean branch lengths of the clade of predicted viral genes vs. mean branch lengths of the clade of reference sequences). Second, to test the relatedness between the reconstructed viral sequences and those in the viral reference database (length of the branch leading to the clade of viral genes vs. mean branch lengths of the clade of reference sequences).

When viral genes were not monophyletic, mean branch lengths within both types of clades were calculated separately using the `drop.tip` function. The length of the branch leading to the clade of genes could not be measured in those cases. Multiple comparisons were performed using the Dunn test with the Benjamini and Hochberg (BH) correction.

## Results

We reconstructed viral genes from High Arctic soil and lake sediments using paired meta-transcriptomic (RNA, *n* = 6) and metagenomic (DNA, *n* = 8) datasets (Figure 1). Assembly yielded a median of 504,778 contigs per sample; RNA and DNA contigs were pooled separately. Among classified contigs, 99% were bacterial, archaeal, or eukaryotic and removed, leaving unclassified contigs and other taxa for viral scanning. Only 0.19% (RNA) and 0.45% (DNA) of sequences were identified as putatively viral; non-proviral sequences with ≥1 viral gene were retained for protein-coding gene prediction.

A reference database of 169.5 million viral proteins from RefSeq v230, GenBank v266, and IMG/VR v4.1 was clustered into 1,910,901 HMM profiles. Our predicted genes were searched against this database, yielding **C**lusters **o**f **P**utatively **H**omologous **Se**quences (CoPHSe) with ≥3 predicted genes and ≥1 reference (Table 1). We recovered 41,883 RNA and 403,266 DNA CoPHSe at *E*-value 10^−15^, and 32,637 RNA and 330,876 DNA CoPHSe at 10^−20^.

**Table 1.** Number of CoPHSe per type and *E*-value.

| Type | 10<sup>-15</sup> |  | 10<sup>-20</sup> |  |
| --- | --- | --- | --- | --- |
|  | RNA<br>N (%) | DNA<br>N (%) | RNA<br>N (%) | DNA<br>N (%) |
| Non-phage | 6,041 (14.4) | 6,716 (1.7) | 5,578 (17.1) | 5,593 (1.7) |
| Phage | 4,523 (10.8) | 6,788 (1.7) | 4,156 (12.7) | 5,649 (1.7) |
| Unknown | 29,082 (69.4) | 354,791 (87.9) | 21,317 (65.3) | 290,391 (87.7) |
| Non-phage, Unknown | 425 (1.0) | 1,613 (0.4) | 398 (1.2) | 1,260 (0.4) |
| Phage, Unknown | 1,812 (4.3) | 33,358 (8.3) | 1,188 (3.6) | 27,983 (8.5) |
| **Total** | **41,883 (100.0)** | **403,266 (100.0)** | **32,637 (100.0)** | **330,876 (100.0)** |

Most CoPHSe (65–69% RNA; 88% DNA) were “unknown.” Phage and non-phage CoPHSe were more abundant in RNA (11–13% phage; 14–17% non-phage) than DNA (2% each), with minor fractions classified as “phage + unknown” (4% RNA; 8–9% DNA).

### High Arctic viruses show greater diversification

To assess diversification independent of taxonomy, we compared mean branch lengths within reconstructed gene clades vs. reference clades across phylogenetic trees (Figure S2). At the stringent *E*-value 10^−20^, non-phage RNA/DNA genes exhibited median branch lengths of 0.24 and 0.26 substitutions/site (sub/site), respectively, compared to 0.0026 and 0.0046 sub/site for references (Figure 2). Phage and unknown categories showed similar patterns: references neared 0 (≤0.0044 sub/site) while genes ranged from 0.13– 0.29 sub/site. All differences were significant (*p <* 0.001; Supplementary Table 1; median 63-fold increase). Results held for *E*-value 10^−15^ (Supplementary Figure 3; Supplementary Table 2) and mixed categories (Supplementary Figure 4; Table S3).

**Figure 2.**
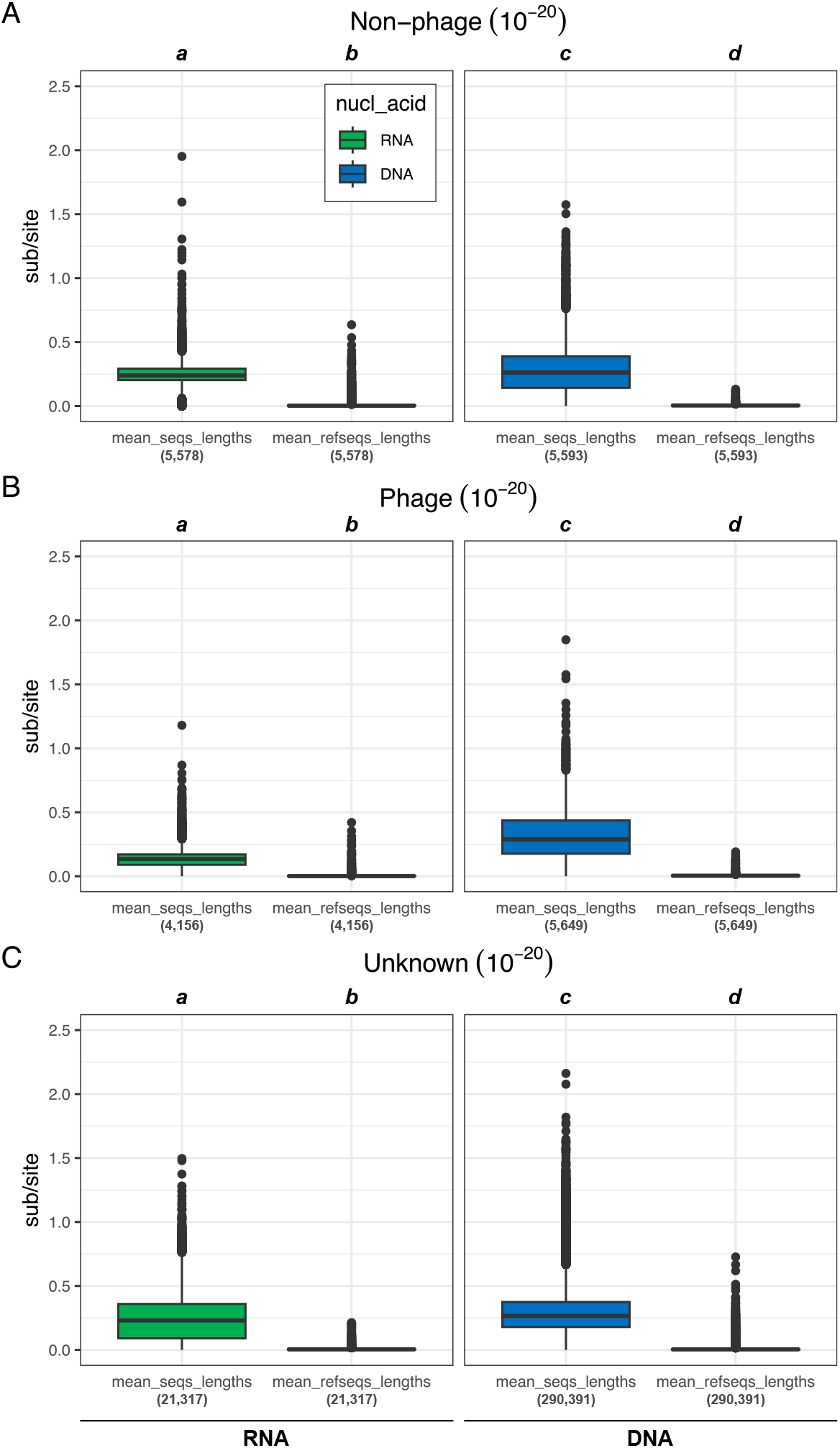
Comparative mean branch length among CoPHSe (“mean_seqs_lengths”) vs. mean branch length among reference sequences (“mean_refseqs_length”). Diversification was assessed in expected numbers of substitutions per site (sub/site). CoPHSe were obtained using the E-value threshold of 10^-20^ (see Supplementary Figure 3 for 10^-15^). (A) Non-phage, (B) Phage, and (C) Unknown RNA (green) and DNA (blue) CoPHSe. For each type of CoPHSe, pairwise multiple comparisons were performed using the Dunn test with the BH correction (*α* = 0.05, Supplementary Table 1). Significant results are marked with letters from *a* to *d*. Numbers of reconstructed trees are shown below each column in parentheses.

### High Arctic viruses are highly divergent from known viruses

We next quantified evolutionary distance by comparing the branch length to reconstructed gene clades against reference clade means (Figure S2). These connecting branches spanned 0.56–1.10 sub/site across non-phage, phage, and unknown CoPHSe (*E*-value 10^−20^; Figure 3), significantly exceeding reference diversification (*p <* 0.001; Supplementary Table 4). Equivalent patterns emerged at *E*-value 10^−15^ (Supplementary Figure 5; Supplementary Table 5) and for mixed categories (Supplementary Figure 6; Supplementary Table 6).

These findings indicate that High Arctic viral genes have diversified substantially more than, and are far more divergent from, sequences in public databases, suggesting distinct evolutionary trajectories.

**Figure 3.**
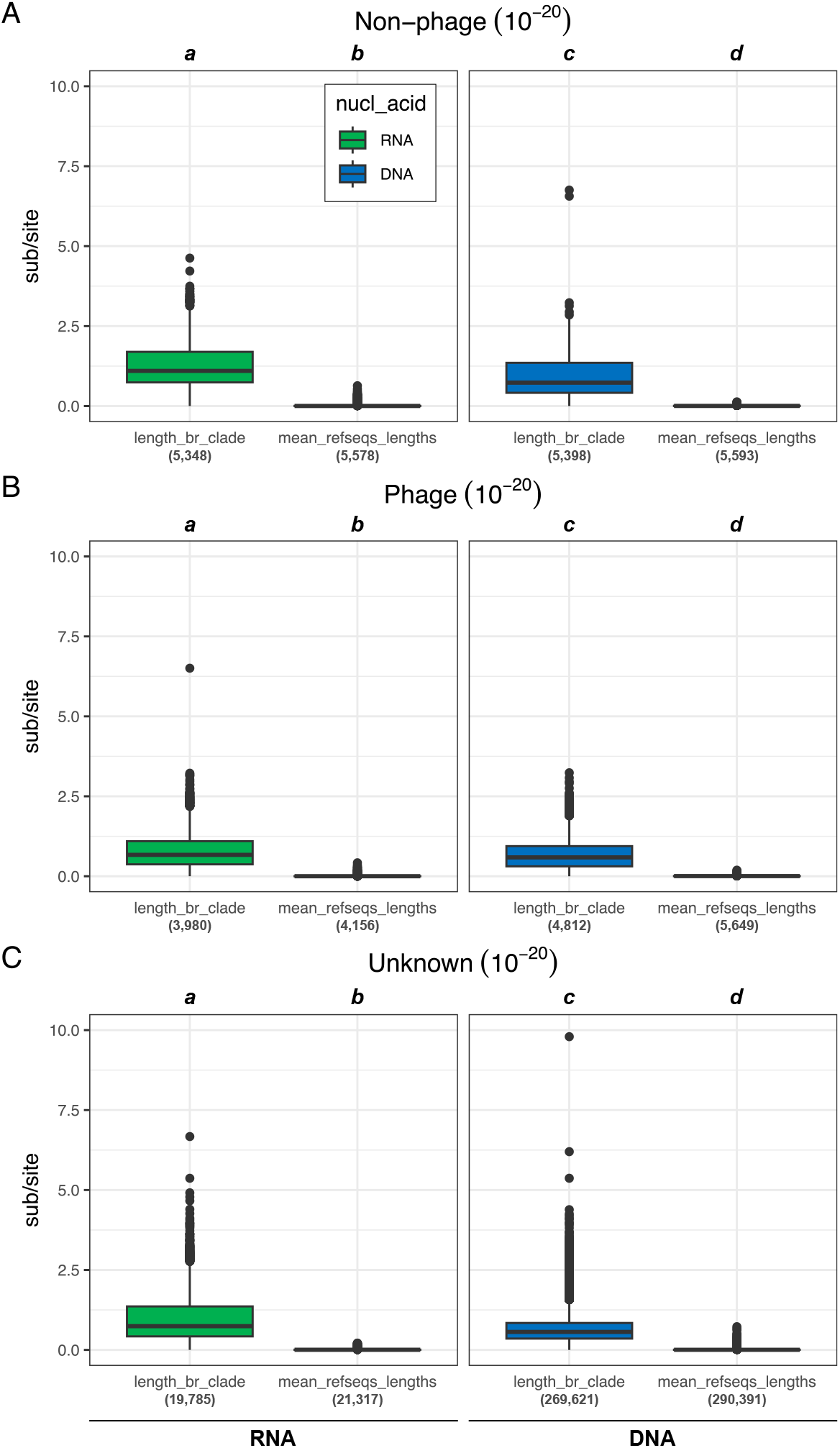
Comparative length of branches leading to CoPHSe (“length_br_clade”) vs. mean branch length among reference sequences (“mean_refseqs_length”). Diversification was assessed in expected numbers of substitutions per site. CoPHSe were obtained using the E-value threshold of 10^-20^ (see Supplementary Figure 5 for 10^-15^). (A) Non-phage, (B) Phage, and (C) Unknown RNA (green) and DNA (blue) CoPHSe. For each type of CoPHSe, pairwise multiple comparisons were performed using the Dunn test with the BH correction (*α* = 0.05, Supplementary Table 4). Significant results are marked with letters from a to d. Numbers of reconstructed trees are shown below each column in parentheses.

## Discussion

High Arctic viruses from Lake Hazen sediments and soils are predominantly novel, with most genes clustering as unknown or distantly related to known sequences. Phages dominated our Clusters of Putatively Homologous Sequences (CoPHSe), consistent with prior work at this site (Lemieux et al., 2022) and other Arctic studies (Dávila-Ramos et al., 2019; Emerson et al., 2018; Zhong et al., 2020; Wang et al., 2022; Allen et al., 2017; Labbé et al., 2020; Pettersson et al., 2025). This phage prevalence likely reflects high bacterial densities in these sediments (Lemieux et al., 2022; Simon, Wiezer, Strittmatter, & Daniel, 2009; Suttner et al., 2020), providing abundant hosts for infection and replication (Anesio, Mindl, Laybourn-Parry, Hodson, & Sattler, 2007).

Our HMM-based phylogenetic clustering approach overcomes limitations of traditional similarity searches by detecting remote homologs among highly divergent sequences. However, reliance on protein-coding gene predictions may underestimate diversity from short contigs or non-coding elements, while assembly biases could favor abundant phages (Skewes-Cox, Sharpton, Pollard, & DeRisi, 2014).

Critically, High Arctic viral genes exhibited substantially greater diversification (median 63-fold higher branch lengths) and divergence (0.56–1.10 sub/site connecting branches) than public database references across all categories. Several non-mutually exclusive mechanisms may explain this pattern. First, isolation by distance and small local effective population sizes could drive neutral drift, with limited gene flow from lower latitudes exacerbating divergence (Lemieux et al., 2022; Zohdy, Schwartz, & Oaks, 2019). Second, adaptation to extreme conditions (e.g., low temperatures, desiccation, freeze-thaw cycles) may accelerate molecular evolution (Gil et al., 2021). Third, the extreme undersampling of polar viromes (Yau & Seth-Pasricha, 2019) means even “known” Arctic viruses remain uncharacterized, as evidenced by frequent unclassified sequences even at superkingdom level (Aguirre de Cárcer et al., 2015; Emerson et al., 2018; Zhong et al., 2020; Wang et al., 2022; Kim et al., 2024; Angly et al., 2006; Demina et al., 2025; Pettersson et al., 2025; Pratama et al., 2025).

As climate warming fragments Arctic habitats and boosts meltwater inputs to Lake Hazen, northward host range shifts could facilitate viral spillover, amplifying pandemic risks in sparsely monitored regions. These findings reveal a highly distinct High Arctic virome that has evolved under unique selective pressures. Future work integrating long-read sequencing and host-range predictions will clarify these dynamics, informing polar microbial responses to global change.

## Acknowledgements

We would like to thank the Digital Research Alliance of Canada (DRAC), formerly known as Compute Canada, for providing us with compute time on their servers.

## Author contributions

Conceptualization: AL, AJP, SAB. Methodology: AL, AJP, SAB. Investigation: AL, SAB. Data curation: AL. Formal analysis: AL, SAB. Visualization: AL, SAB. Writing: original draft: AL, SAB; review & editing: AL, AJP, SAB. Supervision: AJP, SAB.

## Supplementary data

Supplementary data is available at *VEVOLU* online.

## Data availability

The raw data used in this study can be found at https://www.ncbi.nlm.nih.gov/bioproject/PRJNA746497/ and https://www.ncbi.nlm.nih.gov/bioproject/556841. The code developed for this work is available from https://github.com/sarisbro/data.

