## Supplementary material for "The world’s largest High Arctic lake is dominated by uncharacterized, genetically highly diverse phages": SI file

#### Supplementary figures

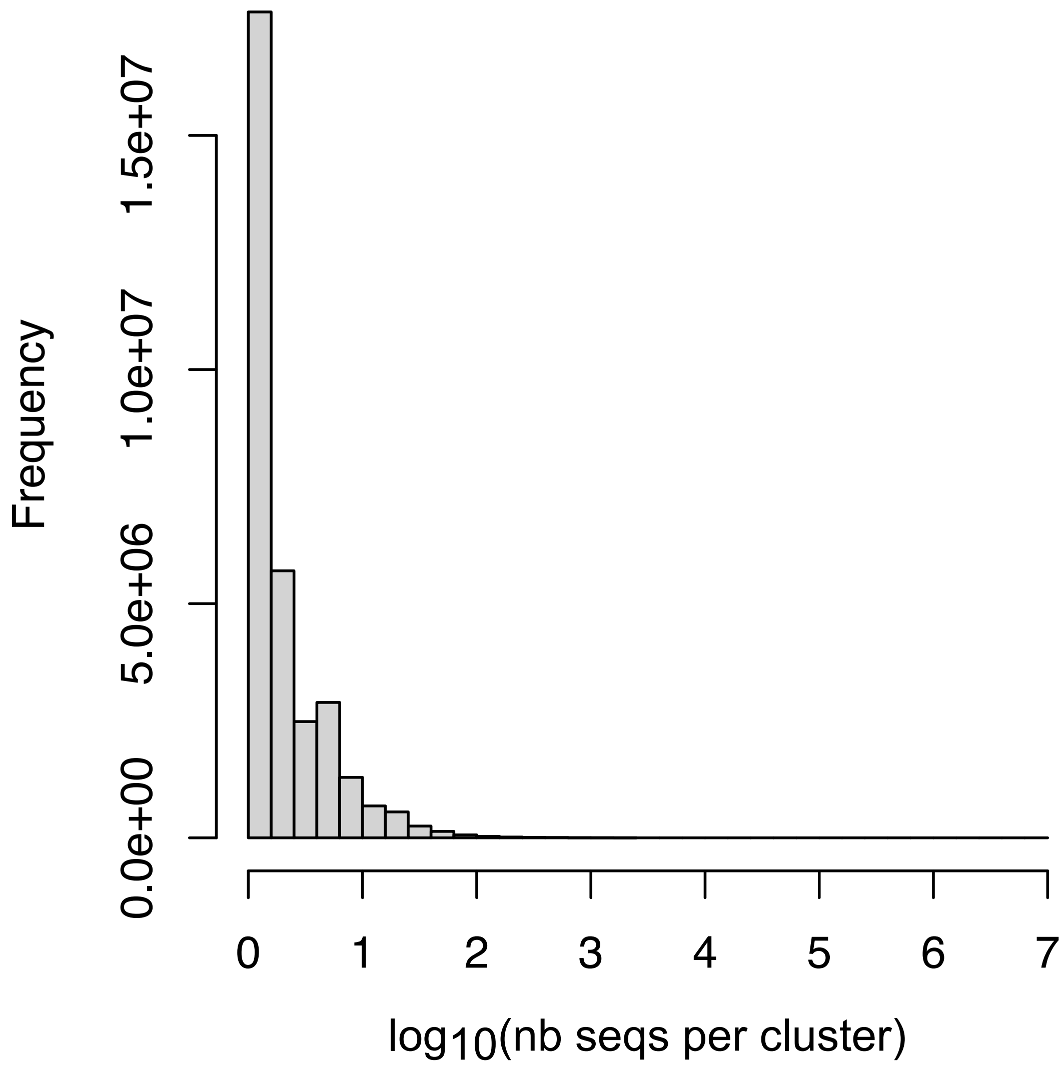


**Supplementary Figure 1. Number of sequences per viral cluster.** The number of sequences is on a $\log_{10}$ scale.

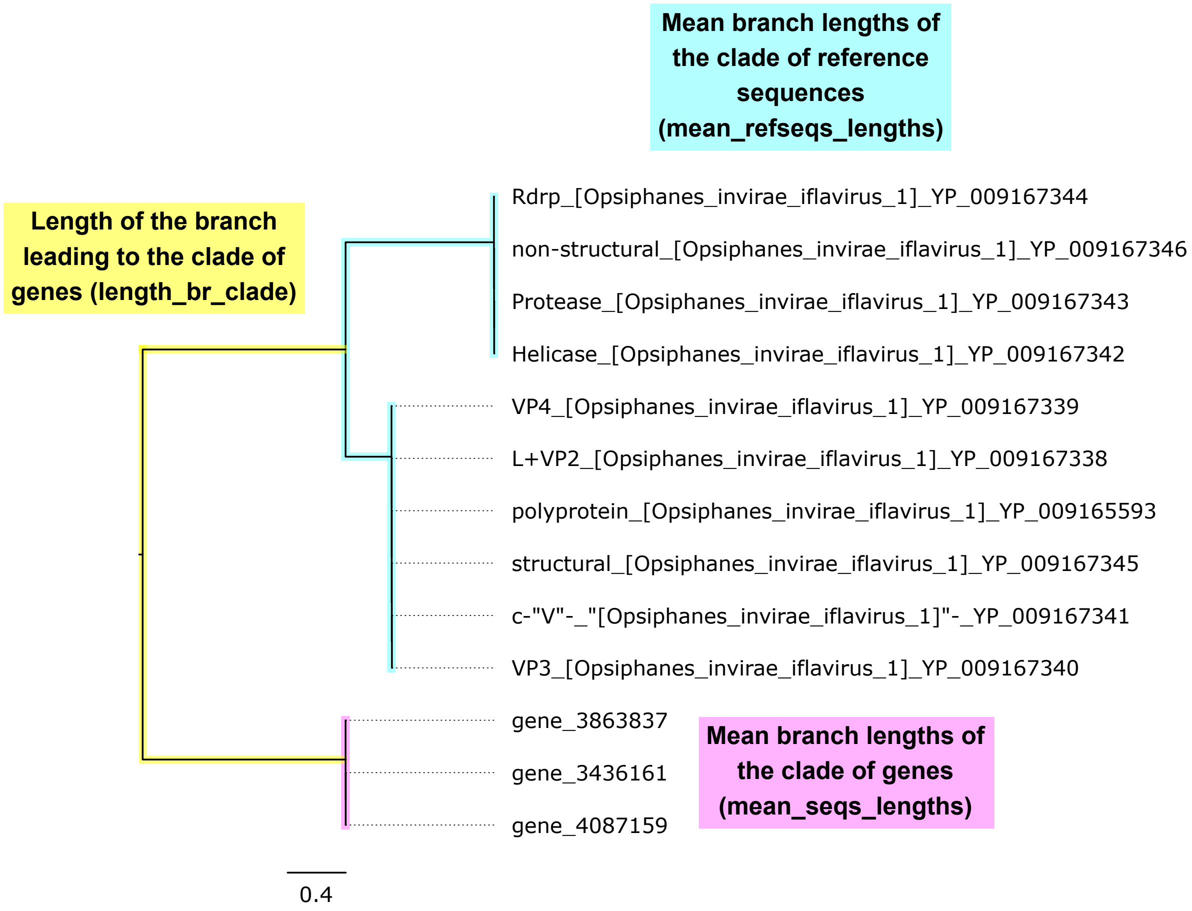


**Supplementary Figure 2.** **Assessment of phylogeny.** Phylogenetic trees were first rerooted to put genes into a single clade. The phylogenies were then assessed in three ways: with (i) the length of the branch leading to the clade of genes ("length_br_clade"; yellow), (ii) the mean length of branches within the clade of genes ("mean_seqs_lengths"; pink), and (iii) the mean length of branches within the clade of reference sequences ("mean_refseqs_lengths"; blue). In cases where genes were not monophyletic, the "length_br_clade" was not assessed.

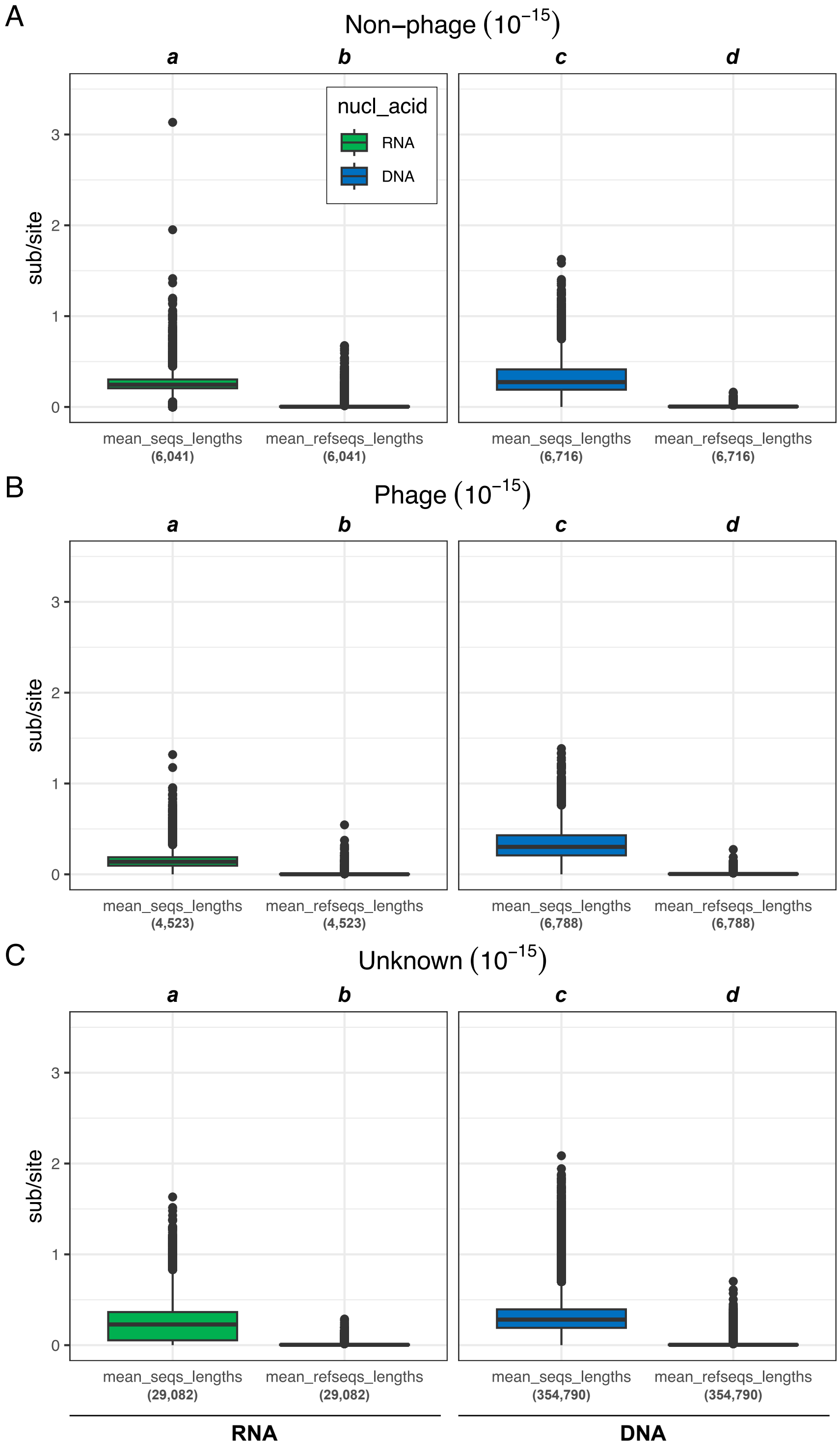


**Supplementary Figure 3.** **Comparative mean branch length among CoPHSe ("mean_seqs_lengths") vs. mean branch length among reference sequences ("mean_seqs_length").** Diversification was assessed in expected numbers of substitutions per site (sub/site). CoPHSe were obtained using the E-value threshold of $10^{-15}$ (see Figure 2 for $10^{-20}$). (A) Non-phage, (B) Phage, and (C) Unknown RNA (green) and DNA (blue) CoPHSe. For each type of CoPHSe, pairwise multiple comparisons were performed using the Dunn test with the BH correction ($\alpha$ = 0.05, Supplementary Table 2). Significant results are marked with letters from *a* to *d*. Numbers of reconstructed trees are shown below each column in parentheses.

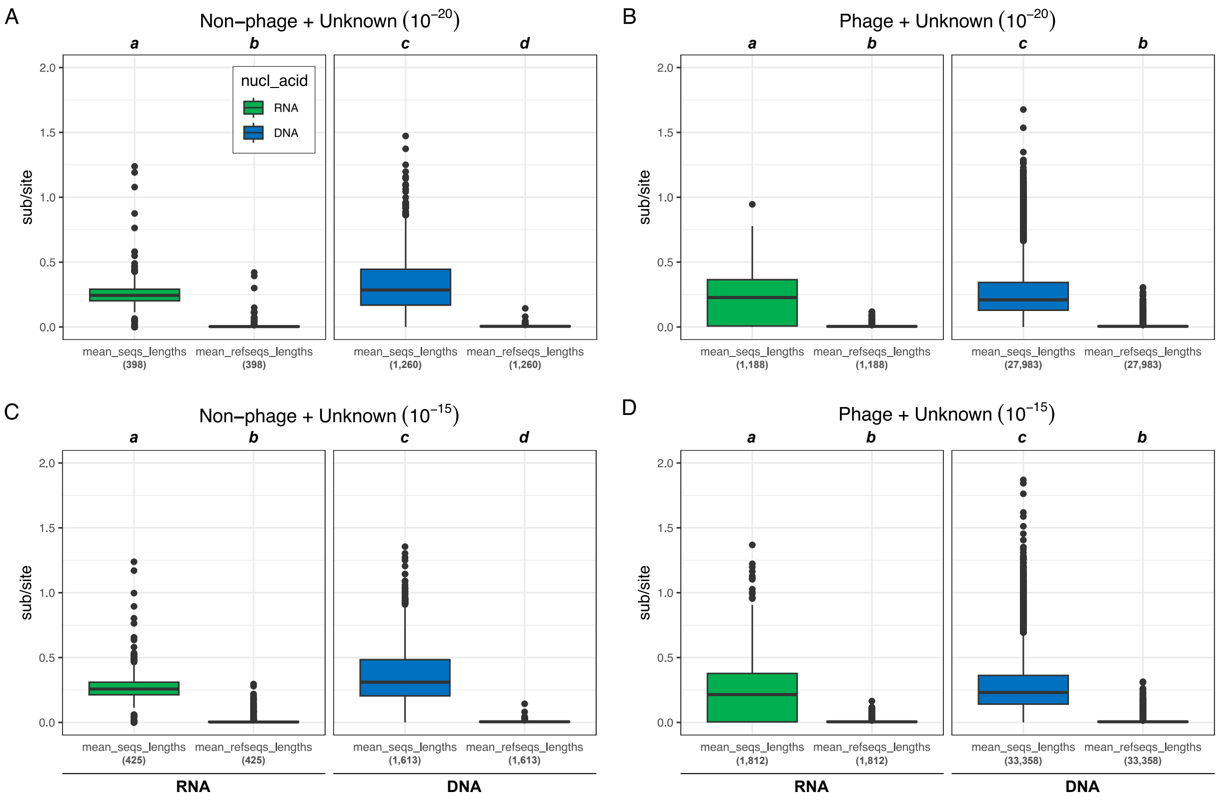


**Supplementary Figure 4. Comparative mean branch length among CoPHSe ("mean_seqs_lengths") vs. mean branch length among reference sequences ("mean_seqs_length").** Diversification was assessed in expected numbers of substitutions per site (sub/site). CoPHSe were obtained using the E-value thresholds of (A, B) $10^{-20}$ and (C, D) $10^{-15}$. (A, C) Non-phage + Unknown and (B, D) Phage + Unknown RNA (green) and DNA (blue) CoPHSe. For each type of CoPHSe, pairwise multiple comparisons were performed using the Dunn test with the BH correction ($\alpha = 0.05$, Supplementary Table 3). Significant results are marked with letters from *a* to *d*. Numbers of reconstructed trees are shown below each column in parentheses.

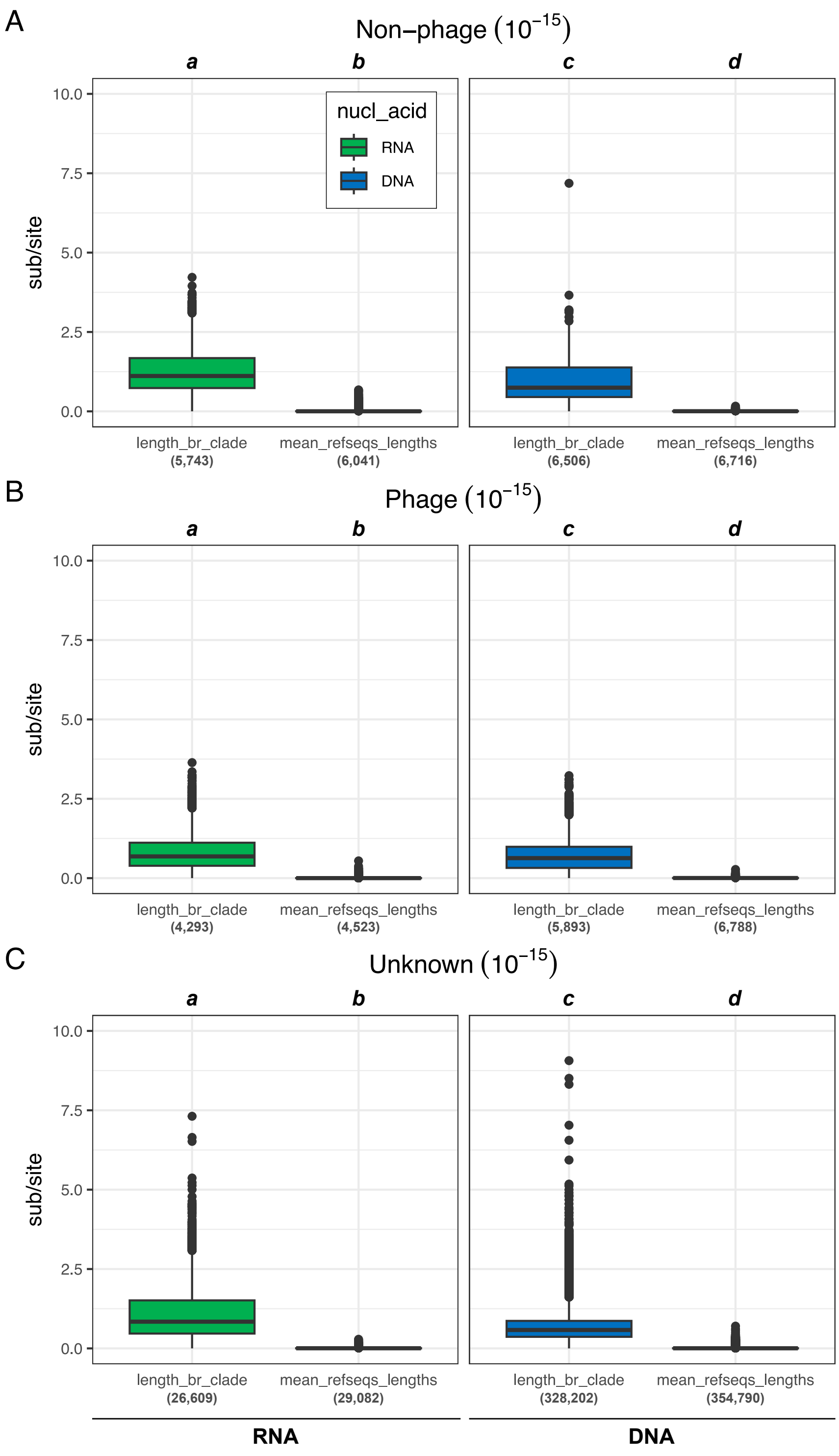


**Supplementary Figure 5. Comparative length of branches leading to CoPHSe ("length_br_clade") vs. mean branch length among reference sequences ("mean_refseqs_length").** Diversification was assessed in expected numbers of substitutions per site. CoPHSe were obtained using the E-value threshold of $10^{-15}$ (see Figure 3 for $10^{-20}$). (A) Non-phage, (B) Phage, and (C) Unknown RNA (green) and DNA (blue) CoPHSe. For each type of CoPHSe, pairwise multiple comparisons were performed using the Dunn test with the BH correction ($\alpha = 0.05$, Supplementary Table 5). Significant results are marked with letters from a to d. Numbers of reconstructed trees are shown below each column in parentheses.

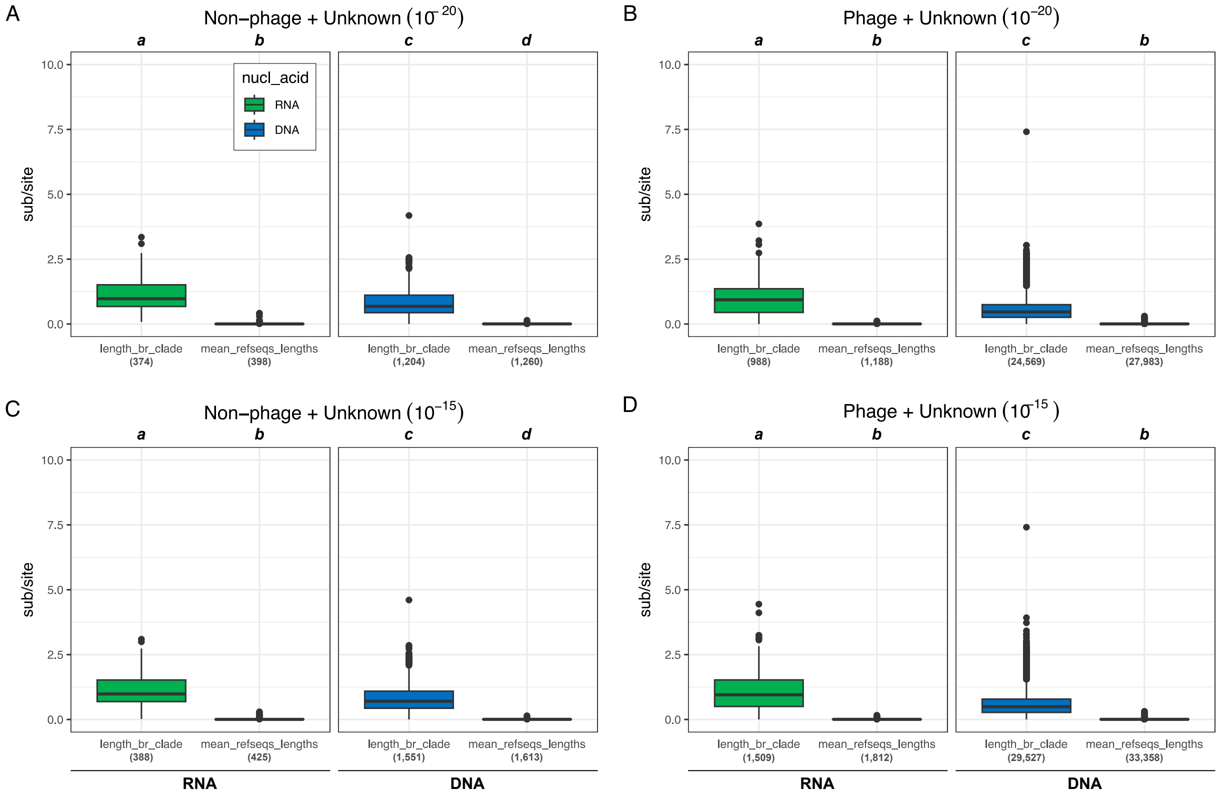


**Supplementary Figure 6. Comparative length of branches leading to CoPHSe ("length_br_clade") vs. mean branch length among reference sequences ("mean_seqs_length").** Diversification was assessed in expected numbers of substitutions per site (sub/site). CoPHSe were obtained using the E-value thresholds of (A, B) $10^{-20}$ and (C, D) $10^{-15}$. (A, C) Non-phage + Unknown and (B, D) Phage + Unknown RNA (green) and DNA (blue) CoPHSe. For each type of CoPHSe, pairwise multiple comparisons were performed using the Dunn test with the BH correction ($\alpha = 0.05$, Supplementary Table 6). Significant results are marked with letters from *a* to *d*. Numbers of reconstructed trees are shown below each column in parentheses.

### Supplementary tables

**Supplementary Table 1.** Pairwise multiple comparisons between the "mean_seqs_length" and the "mean_refseqs_length" metrics in non-phage, phage, and unknown RNA and DNA CoPHSe obtained with the $E$-value threshold $10^{-20}$. P-values were computed from the Dunn test with the BH correction. Significant results ($\alpha$ = 0.05) are in bold.

| | | RNA | | DNA | |
|---|---|---|---|---|---|
| | | mean_seqs_lengths | mean_refseqs_lengths | mean_seqs_lengths | mean_refseqs_lengths |
| Non-phage CoPHSe ($10^{-20}$) | | | | | |
| RNA | mean_seqs_lengths | – | **<0.001** | **<0.001** | **<0.001** |
| | mean_refseqs_lengths | – | – | **<0.001** | **<0.001** |
| DNA | mean_seqs_lengths | – | – | – | **<0.001** |
| | mean_refseqs_lengths | – | – | – | – |
| Phage CoPHSe ($10^{-20}$) | | | | | |
| RNA | mean_seqs_lengths | – | **<0.001** | **<0.001** | **<0.001** |
| | mean_refseqs_lengths | – | – | **<0.001** | **<0.001** |
| DNA | mean_seqs_lengths | – | – | – | **<0.001** |
| | mean_refseqs_lengths | – | – | – | – |
| Unknown CoPHSe ($10^{-20}$) | | | | | |
| RNA | mean_seqs_lengths | – | **<0.001** | **<0.001** | **<0.001** |
| | mean_refseqs_lengths | – | – | **<0.001** | **<0.001** |
| DNA | mean_seqs_lengths | – | – | – | **<0.001** |
| | mean_refseqs_lengths | – | – | – | – |

**Supplementary Table 2.** Pairwise multiple comparisons between the "mean_seqs_length" and the "mean_refseqs_length" metrics in non-phage, phage, and unknown RNA and DNA CoPHSe obtained with the $E$-value threshold $10^{-15}$. P-values were computed from the Dunn test with the BH correction. Significant results ($\alpha = 0.05$) are in bold.

| | | RNA | | DNA | |
|---|---|---|---|---|---|
| | | mean_seqs_lengths | mean_refseqs_lengths | mean_seqs_lengths | mean_refseqs_lengths |
| Non-phage CoPHSe ($10^{-15}$) | | | | | |
| RNA | mean_seqs_lengths | – | **<0.001** | **<0.001** | **<0.001** |
| | mean_refseqs_lengths | – | – | **<0.001** | **<0.001** |
| DNA | mean_seqs_lengths | – | – | – | **<0.001** |
| | mean_refseqs_lengths | – | – | – | – |
| Phage CoPHSe ($10^{-15}$) | | | | | |
| RNA | mean_seqs_lengths | – | **<0.001** | **<0.001** | **<0.001** |
| | mean_refseqs_lengths | – | – | **<0.001** | **<0.001** |
| DNA | mean_seqs_lengths | – | – | – | **<0.001** |
| | mean_refseqs_lengths | – | – | – | – |
| Unknown CoPHSe ($10^{-15}$) | | | | | |
| RNA | mean_seqs_lengths | – | **<0.001** | **<0.001** | **<0.001** |
| | mean_refseqs_lengths | – | – | **<0.001** | **<0.001** |
| DNA | mean_seqs_lengths | – | – | – | **<0.001** |
| | mean_refseqs_lengths | – | – | – | – |

**Supplementary Table 3.** Pairwise multiple comparisons between the "mean_seqs_length" and the "mean_refseqs_length" metrics in non-phage + unknown and phage + unknown RNA and DNA CoPHSe obtained with the $E$-value thresholds $10^{-20}$ and $10^{-15}$. P-values were computed from the Dunn test with the BH correction. Significant results ($\alpha = 0.05$) are in bold.

| | | RNA | | DNA | |
|---|---|---|---|---|---|
| | | mean_seqs_lengths | mean_refseqs_lengths | mean_seqs_lengths | mean_refseqs_lengths |
| Non-phage + Unknown CoPHSe ($10^{-20}$) | | | | | |
| RNA | mean_seqs_lengths | – | **<0.001** | **<0.001** | **<0.001** |
| | mean_refseqs_lengths | – | – | **<0.001** | **0.007** |
| DNA | mean_seqs_lengths | – | – | – | **<0.001** |
| | mean_refseqs_lengths | – | – | – | – |
| Phage + Unknown CoPHSe ($10^{-20}$) | | | | | |
| RNA | mean_seqs_lengths | – | **<0.001** | **<0.001** | **<0.001** |
| | mean_refseqs_lengths | – | – | **<0.001** | 0.371 |
| DNA | mean_seqs_lengths | – | – | – | **<0.001** |
| | mean_refseqs_lengths | – | – | – | – |
| Non-phage + Unknown CoPHSe ($10^{-15}$) | | | | | |
| RNA | mean_seqs_lengths | – | **<0.001** | **<0.001** | **<0.001** |
| | mean_refseqs_lengths | – | – | **<0.001** | **0.021** |
| DNA | mean_seqs_lengths | – | – | – | **<0.001** |
| | mean_refseqs_lengths | – | – | – | – |
| Phage + Unknown CoPHSe ($10^{-15}$) | | | | | |
| RNA | mean_seqs_lengths | – | **<0.001** | **<0.001** | **<0.001** |
| | mean_refseqs_lengths | – | – | **<0.001** | 0.072 |
| DNA | mean_seqs_lengths | – | – | – | **<0.001** |
| | mean_refseqs_lengths | – | – | – | – |

**Supplementary Table 4.** Pairwise multiple comparisons between the "length_br_clade" and the "mean_refseqs_length" metrics in non-phage, phage, and unknown RNA and DNA CoPHSe obtained with the $E$-value threshold $10^{-20}$. P-values were computed from the Dunn test with the BH correction. Significant results ($\alpha$ = 0.05) are in bold.

| | | RNA | | DNA | |
|---|---|---|---|---|---|
| | | length_br_clade | mean_refseqs_lengths | length_br_clade | mean_refseqs_lengths |
| Non-phage CoPHSe ($10^{-20}$) | | | | | |
| RNA | length_br_clade | – | **<0.001** | **<0.001** | **<0.001** |
| | mean_refseqs_lengths | – | – | **<0.001** | **<0.001** |
| DNA | length_br_clade | – | – | – | **<0.001** |
| | mean_refseqs_lengths | – | – | – | – |
| Phage CoPHSe ($10^{-20}$) | | | | | |
| RNA | length_br_clade | – | **<0.001** | **<0.001** | **<0.001** |
| | mean_refseqs_lengths | – | – | **<0.001** | **<0.001** |
| DNA | length_br_clade | – | – | – | **<0.001** |
| | mean_refseqs_lengths | – | – | – | – |
| Unknown CoPHSe ($10^{-20}$) | | | | | |
| RNA | length_br_clade | – | **<0.001** | **<0.001** | **<0.001** |
| | mean_refseqs_lengths | – | – | **<0.001** | **<0.001** |
| DNA | length_br_clade | – | – | – | **<0.001** |
| | mean_refseqs_lengths | – | – | – | – |

**Supplementary Table 5.** Pairwise multiple comparisons between the "length_br_clade" and the "mean_refseqs_length" metrics in non-phage, phage, and unknown RNA and DNA CoPHSe obtained with the $E$-value threshold $10^{-15}$. P-values were computed from the Dunn test with the BH correction. Significant results ($\alpha$ = 0.05) are in bold.

| | | RNA | | DNA | |
|---|---|---|---|---|---|
| | | length_br_clade | mean_refseqs_lengths | length_br_clade | mean_refseqs_lengths |
| Non-phage CoPHSe ($10^{-15}$) | | | | | |
| RNA | length_br_clade | – | **<0.001** | **<0.001** | **<0.001** |
| | mean_refseqs_lengths | – | – | **<0.001** | **<0.001** |
| DNA | length_br_clade | – | – | – | **<0.001** |
| | mean_refseqs_lengths | – | – | – | – |
| Phage CoPHSe ($10^{-15}$) | | | | | |
| RNA | length_br_clade | – | **<0.001** | **<0.001** | **<0.001** |
| | mean_refseqs_lengths | – | – | **<0.001** | **<0.001** |
| DNA | length_br_clade | – | – | – | **<0.001** |
| | mean_refseqs_lengths | – | – | – | – |
| Unknown CoPHSe ($10^{-15}$) | | | | | |
| RNA | length_br_clade | – | **<0.001** | **<0.001** | **<0.001** |
| | mean_refseqs_lengths | – | – | **<0.001** | **<0.001** |
| DNA | length_br_clade | – | – | – | **<0.001** |
| | mean_refseqs_lengths | – | – | – | – |

**Supplementary Table 6.** Pairwise multiple comparisons between the "length_br_clade" and the "mean_refseqs_length" metrics in non-phage + unknown and phage + unknown RNA and DNA CoPHSe obtained with the $E$-value thresholds $10^{-20}$ and $10^{-15}$. P-values were computed from the Dunn test with the BH correction. Significant results ($\alpha = 0.05$) are in bold.

| | | RNA | | DNA | |
|---|---|---|---|---|---|
| | | length_br_clade | mean_refseqs_lengths | length_br_clade | mean_refseqs_lengths |
| Non-phage + Unknown CoPHSe (10-20) | | | | | |
| RNA | length_br_clade | – | **<0.001** | **<0.001** | **<0.001** |
| | mean_refseqs_lengths | – | – | **<0.001** | **0.004** |
| DNA | length_br_clade | – | – | – | **<0.001** |
| | mean_refseqs_lengths | – | – | – | – |
| Phage + Unknown CoPHSe (10-20) | | | | | |
| RNA | length_br_clade | – | **<0.001** | **<0.001** | **<0.001** |
| | mean_refseqs_lengths | – | – | **<0.001** | 0.060 |
| DNA | length_br_clade | – | – | – | **<0.001** |
| | mean_refseqs_lengths | – | – | – | – |
| Non-phage + Unknown CoPHSe (10-15) | | | | | |
| RNA | length_br_clade | – | **<0.001** | **<0.001** | **<0.001** |
| | mean_refseqs_lengths | – | – | **<0.001** | **0.013** |
| DNA | length_br_clade | – | – | – | **<0.001** |
| | mean_refseqs_lengths | – | – | – | – |
| Phage + Unknown CoPHSe (10-15) | | | | | |
| RNA | length_br_clade | – | **<0.001** | **<0.001** | **<0.001** |
| | mean_refseqs_lengths | – | – | **<0.001** | 0.348 |
| DNA | length_br_clade | – | – | – | **<0.001** |
| | mean_refseqs_lengths | – | – | – | – |
